# Favipiravir-resistant influenza A virus shows potential for transmission

**DOI:** 10.1101/2020.09.01.277343

**Authors:** Daniel H. Goldhill, Ada Yan, Rebecca Frise, Jie Zhou, Jennifer Shelley, Ana Gallego Cortés, Shahjahan Miah, Omolola Akinbami, Monica Galiano, Maria Zambon, Angie Lackenby, Wendy S. Barclay

## Abstract

Favipiravir is a nucleoside analogue which has been licensed to treat influenza in the event of a new pandemic. We previously described a favipiravir resistant influenza A virus generated by in vitro passage in presence of drug with two mutations: K229R in PB1, which conferred resistance at a cost to polymerase activity, and P653L in PA, which compensated for the cost of polymerase activity. However, the clinical relevance of these mutations is unclear as the mutations have not been found in natural isolates and it is unknown whether viruses harbouring these mutations would replicate or transmit in vivo. Here, we infected ferrets with a mix of wild type p(H1N1) 2009 and corresponding favipiravir-resistant virus and tested for replication and transmission in the absence of drug. Favipiravir-resistant virus successfully infected ferrets and was transmitted by both contact transmission and respiratory droplet routes. However, sequencing revealed the mutation that conferred resistance, K229R, decreased in frequency over time within ferrets. Modelling revealed that due to a fitness advantage for the PA P653L mutant, reassortment with the wild-type virus to gain wild-type PB1 segment in vivo resulted in the loss of the PB1 resistance mutation K229R. We demonstrated that this fitness advantage of PA P653L in the background of our starting virus A/England/195/2009 was due to a maladapted PA in first wave isolates from the 2009 pandemic. We show there is no fitness advantage of P653L in more recent pH1N1 influenza A viruses. Therefore, whilst favipiravir-resistant virus can transmit in vivo, the likelihood that the resistance mutation is retained in the absence of drug pressure may vary depending on the genetic background of the starting viral strain.

**Author Summary:** In the event of a new influenza pandemic, drugs will be our first line of defence against the virus. However, drug resistance has proven to be particularly problematic to drugs against influenza. Favipiravir is a novel drug which might be used against influenza virus in the event of a new pandemic. Is resistance likely to be a problem for the use of favipiravir? Our previous work has shown that resistance to favipiravir can be generated in cell culture but we don’t know whether there will be a cost preventing the spread of resistance in whole organisms. Here, we used a mix of wild-type and resistant influenza viruses from early in the 2009 pandemic to test whether viruses resistant to favipiravir could transmit between ferrets. We found that the resistant viruses could transmit but that the resistance mutation was selected against within some ferrets. Using modelling and in vitro experiments, we found that the resistant mutation was selected against in the influenza strain from our experiment but not in more recently evolved strains. Our results show that favipiravir resistant viruses could spread if resistance is generated but the probability will depend on the genetic background of the virus.

## Introduction

Influenza virus is a negative strand virus that causes significant morbidity and mortality worldwide. Like most other RNA viruses, influenza virus has a fast rate of evolution which allows it to evolve in response to immune pressure as well as antiviral drugs(1-3). Antiviral resistance has been a major problem limiting the effectiveness of antiviral drugs against influenza(4, 5). Significant resistance has evolved against the two main classes of antiviral drugs, adamantanes and neuraminidase inhibitors, which have been used clinically against influenza(5-8). Resistance can also evolve to baloxavir, a recently approved drug that inhibits the cap-snatching ability of the polymerase(9). When developing new drugs, it is vital to understand both the ease with which resistance evolves and the likelihood that resistant variants will transmit especially in the absence of drug pressure. This will inform whether a new drug can be effectively used against influenza on a global scale.

Favipiravir is a novel antiviral drug licensed in Japan for the treatment of influenza in the event of a new pandemic(10-12). Favipiravir is a nucleoside analogue which targets the influenza polymerase and acts as a mutagen(13-15). Favipiravir is active against influenza A and B viruses including strains that are resistant to other classes of antiviral drugs(10, 16). Previously, we demonstrated that Influenza A/England/195/2009 (Eng195), an early isolate from the 2009 H1N1 pandemic, could evolve resistance to favipiravir(17). We showed that two mutations, K229R in PB1 and P653L in PA, were needed to evolve resistance(17). K229R provided resistance to favipiravir at a cost to polymerase activity which was compensated by P653L. K229R has not been found in any natural pH1N1(2009) isolates, which is unsurprising as favipiravir has not been widely used to treat influenza cases. The P653L mutation has also not been found in any natural pH1N1(2009) isolates, which may suggest that it does not confer a fitness advantage in vivo. Although the P653L fully compensated for the cost to polymerase activity and virus replication in vitro, it is unknown whether the resistant virus would replicate in vivo or transmit whilst maintaining resistance.

Transmission studies in animal models such as the ferret have revealed whether drug resistant influenza viruses have fitness costs and can help inform about the likelihood of the emergence of resistance(18-20). Early studies with the adamantane, rimantadine, suggested that there was no fitness cost preventing transmission of resistant influenza in a household setting(21) and resistance has indeed become widespread. Oseltamivir resistant H1N1 viruses were originally shown to be unlikely to transmit(22) but around 2007 additional mutations in the N1 neuraminidase emerged that were subsequently shown to affect NA such that the resistance mutation actually conferred a fitness advantage(5) and widespread resistance to oseltamivir ensued(23). More recently, oseltamivir resistant pH1N1(2009)(24) and baloxavir resistant H3N2 influenza A virus have been shown to transmit between ferrets without a fitness cost(25, 26). In humans so far, resistance to these drugs appears mostly limited to treated patients or small outbreaks but these animal transmission studies suggest that continued use of monotherapy may lead to more widespread resistance.

Interestingly, favipiravir resistant chikungunya virus containing the equivalent mutation to K229R in influenza, has been shown to reproduce less efficiently in mosquitos which led to slower transmission(27). This fitness cost in mosquitos was unexpected as there was no difference in fitness between favipiravir resistant virus and wild-type virus in mammalian cell culture(27, 28). In the present study, we infected ferrets with a mixture of wild-type and resistant virus and tested whether resistant virus would transmit or be outcompeted by the wild-type virus. We constructed a simple model to explain the changes in genotype frequencies observed in our experiment. Finally, we compared pH1N1 PA sequences from 2009-2010 to test whether our results were contingent on the genetic background of the virus.

## Results

### Favipiravir resistant virus transmits between ferrets

To test whether favipiravir resistant virus could transmit by direct or indirect contact, we inoculated ferrets with a mix of resistant virus bearing PB1 K229R + PA P653L and the corresponding wild-type virus, Eng195, a prototypical first wave pH1N1 2009 virus. By inoculating with a mix of virus, we could investigate whether there was a detectable fitness difference between the favipiravir resistant and the wild-type virus. We used a low percentage of wild-type virus to maximize the probability of resistant virus transmitting in the event that there was a fitness cost to resistance in the ferrets. Four donor ferrets were inoculated with K229R + P653L and Eng195 viruses in the ratio of 95:5. After 24 hours, each donor ferret was housed with a direct contact sentinel ferret to measure contact transmission. In addition, an indirect contact sentinel animal was housed in a separate adjacent cage to measure airborne transmission.

All 4 donor ferrets were successfully infected and shed virus in the nasal wash with a peak viral titre on day 2 and a secondary peak for most donors on day 4 or 5 as has been seen previously for ferrets infected with this dose of pH1N1 virus(30) (Figure 1). All 4 direct contact sentinels became infected with the first positive nasal washes occurring between days 2-5. 3 of the 4 indirect contact sentinels became infected and their first positive nasal washes occurred between 3-7 days following infection of the donors. The direct contact and indirect contact sentinels had peak viral titres comparable to the donors suggesting that they were robustly infected.

**Figure 1.**
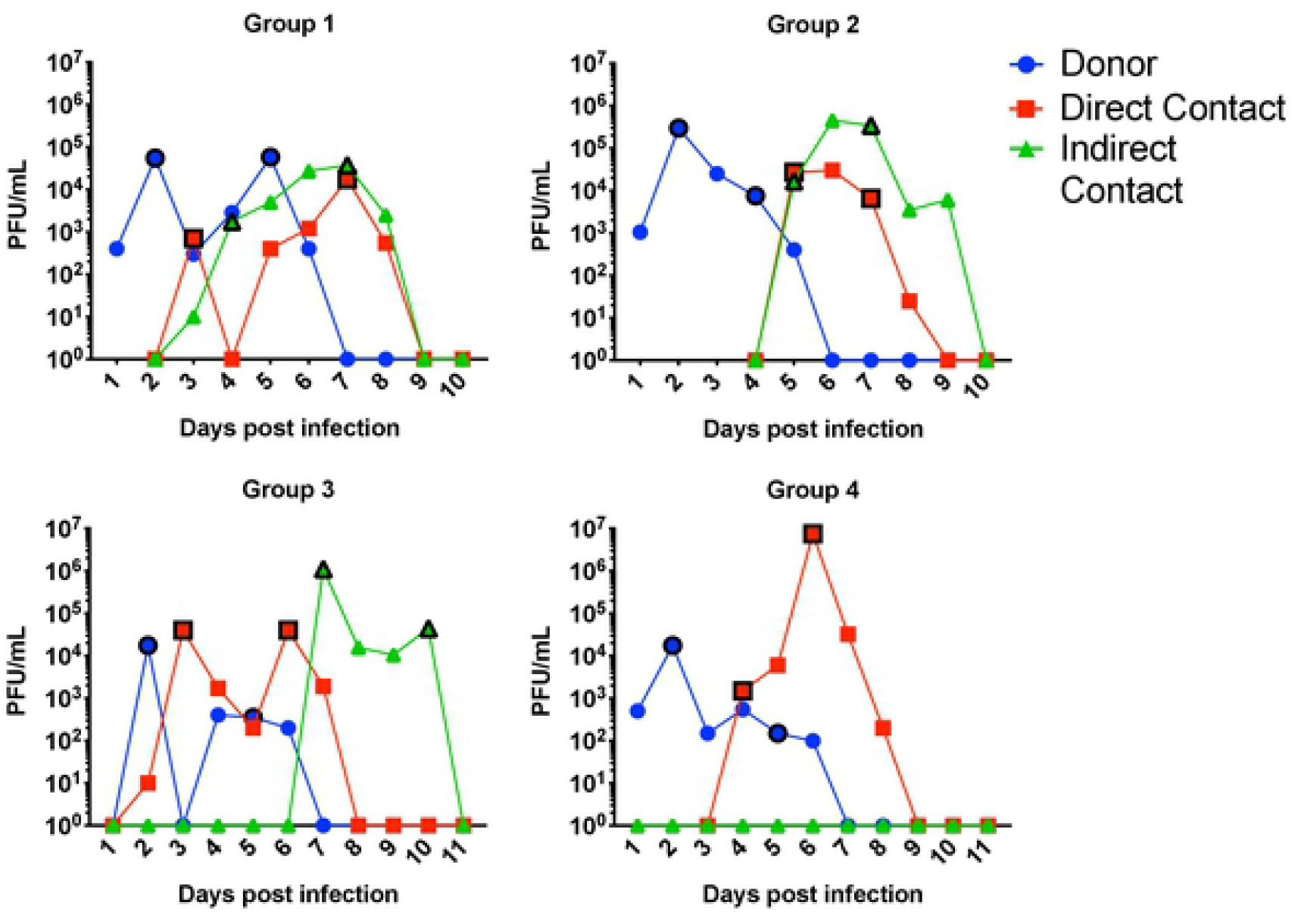
4 donor ferrets were infected with 10^4 PFU of a virus mix of wildtype Eng195 and K229R+P653L. Direct contact and indirect sentinels were exposed from day 1. Ferrets were nasal washed each day and virus infectivity in nasal wash titred by plaque assay. 2 samples were chosen for sequencing from each ferret and are denoted by the black outlined symbols.

Two time points were selected from the daily nasal washes collected from each ferret to sequence virus shed in the nasal washes by both whole genome sequencing and more targeted sequencing of the PB1 and PA segments (Figure 1). Different time points were selected for each individual animal based on their shedding kinetics to give an early snapshot of the diversity of viruses shortly after infection and a later time point to show how viral genotypes change within a host ferret over time. PB1 and PA sequencing revealed that RNA ratio in the inoculum was 95% K229R + P653L and 5% Eng195. Sequencing of virus in the nasal wash showed high levels of both R229 in PB1 and L653 in PA in all infected ferrets indicating that the K229R + P653L virus could productively infect ferrets and was efficiently transmitted both through direct and indirect contact transmission routes (Figure 2). The earliest sample after acquisition of virus in 2 of 4 contact ferrets and 2 of 3 indirect contact ferrets contained 100% K229R + P653L, with no transmission of any wild-type segments.

**Figure 2.**
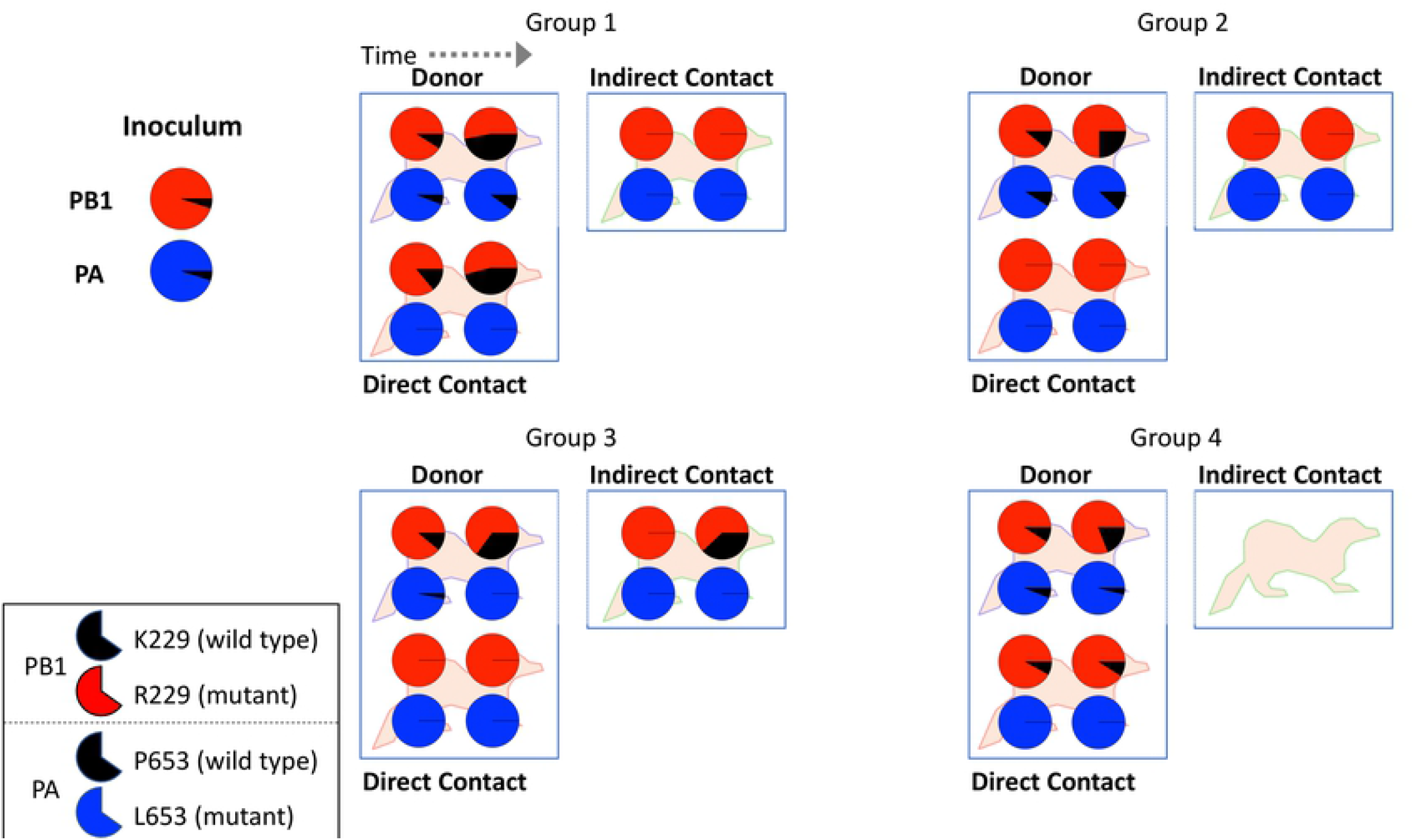
Targeted sequencing of PA and PB1 using NGS showed the percentage of PB1 K229R and PA P653L mutations for donor, contact and indirect contact ferrets. The top pie chart shows the percentage of each genotype for residue 229 in PB1 with the mutant (R229) in red and the wild type (K229) in black. The bottom pie chart shows the percentage of each genotype for residue 653 in PA with the mutant (L653) in blue and the wild type (P653) in black. The inoculum shows 5% K229 and 5% P653. For each infected ferret, two sequenced time points (as described in Figure 1) are shown. The group 4 indirect contact was not infected.

### Whole genome sequencing reveals no additional changes in PB1 and PA

Viruses shed in nasal wash underwent whole genome sequencing to search for additional mutations which might be required for efficient transmission or to further compensate for K229R or P653L. We found no additional mutations in polymerase genes in any of the donor ferrets or the sentinel ferrets occurring above 5%. This confirmed that the K229R + P653L virus was able to productively infect and transmit between ferrets. There were a low number of mutations seen in direct contact and respiratory sentinel ferrets which were likely due to bottlenecking occurring during transmission as some of these mutations were present at low percentages in the inoculum. There was no pattern of repeated mutations across different ferrets which would have indicated positive selection.

### R229 mutation selected against over time within ferrets

Next, we sought to understand how the K229R and P653L mutations might change over time within an individual ferret. In all four donor ferrets, the proportion of the PB1 wild-type amino acid, K229 increased compared to R229 from the inoculum at the earliest time point and further increased at the later time point (Figure 2). From 5% in the inoculum, K229 increased to an average of 10% in donors on day 2 and an average of 32% at the second sequencing time point (day 4 or 5). The largest increase was in donor 1 where K229 increased to 47%. By contrast, the percentage of the PA mutations showed a very different pattern in the donor ferrets, with two ferrets showing a slight increase in the wild-type PA amino acid P653, and two a decrease in P653 on day 4/5. P653 increased slightly on average to 6% on day 2 and 6.5% on day 4/5. Sequencing of nasal wash from sentinel ferrets revealed that P653 never transmitted whereas K229 was found in 2/4 contact sentinels and 1/3 aerosol sentinels. When K229 transmitted to a sentinel ferret, the frequency of K229 increased over time in a manner similar to the donor ferret.

### Modelling shows reassortment coupled with a selective advantage for the P653L mutant drives genotype frequency changes

Influenza virus has a segmented genome and the key mutations in our study are located on discrete RNA segments. Next-generation sequencing does not allow detection of linkage between segments, so it is not always possible to know the exact proportion of genotypes of viruses in mixed samples. In the inoculum, PB1 K229 and PA P653 always occur together in wild-type viruses but within the donor ferrets, the evolutionary trajectory of K229 was decoupled from P653. This could have been either due to reversion of the R229 resistance mutation, which is known to have a fitness cost, or to a fitness advantage of reassortant viruses with the mutated PA segment. To understand what processes could be driving the observed genetic changes in this experiment, we constructed a simple model of virus growth dynamics. As significant frequency changes occurred within donors between day 2 and 5 of our experiment when nasal wash titres showed no increase in viral population size, we modelled a fixed maximum population size for viruses and a fixed number of cells replenished each generation. We noted that previous fitness data we generated in MDCK cells showed a large fitness disadvantage for the K229R mutant, little difference between wild type and the double mutants, and potentially, a slight fitness advantage for the P653L mutant(17). Therefore, for our baseline model, we assigned equal fitness to the wild-type virus and K229R + P653L mutant, a fitness advantage to the P653L mutant and set the fitness of the K229R mutant to 0.01.

Allowing for reassortment coupled with a fitness advantage for the P653L mutant, our model showed an increase in the frequency of K229 and a complete loss of P653 due to the increase in the P653L reassortant (Figure 3a). The increase of frequency in K229 was representative of the dynamics seen within the donor ferrets where K229 increased from 5% to >40% in some ferrets. We tested whether the observed increase of K229 could be driven solely by the fitness cost to the K229R mutant. However, without a fitness advantage to the P653L single mutant, the frequency of K229 did not rise above 5% in our model, the starting frequency in the inoculum (Figure 3b). Next, we tested whether the fitness advantage of the P653L single mutant was sufficient to drive the observed dynamics (Figure 3c). Even without a fitness cost to the K229R mutant, there was still a large increase in the proportion of K229 as in the original model as well as a small proportion of K229R single mutant viruses which did not rise above 5%. Therefore, this model demonstrated that the fitness advantage of the PA P653L single mutant was necessary and sufficient to explain the loss of the R229 PB1 resistance mutation.

**Figure 3.**
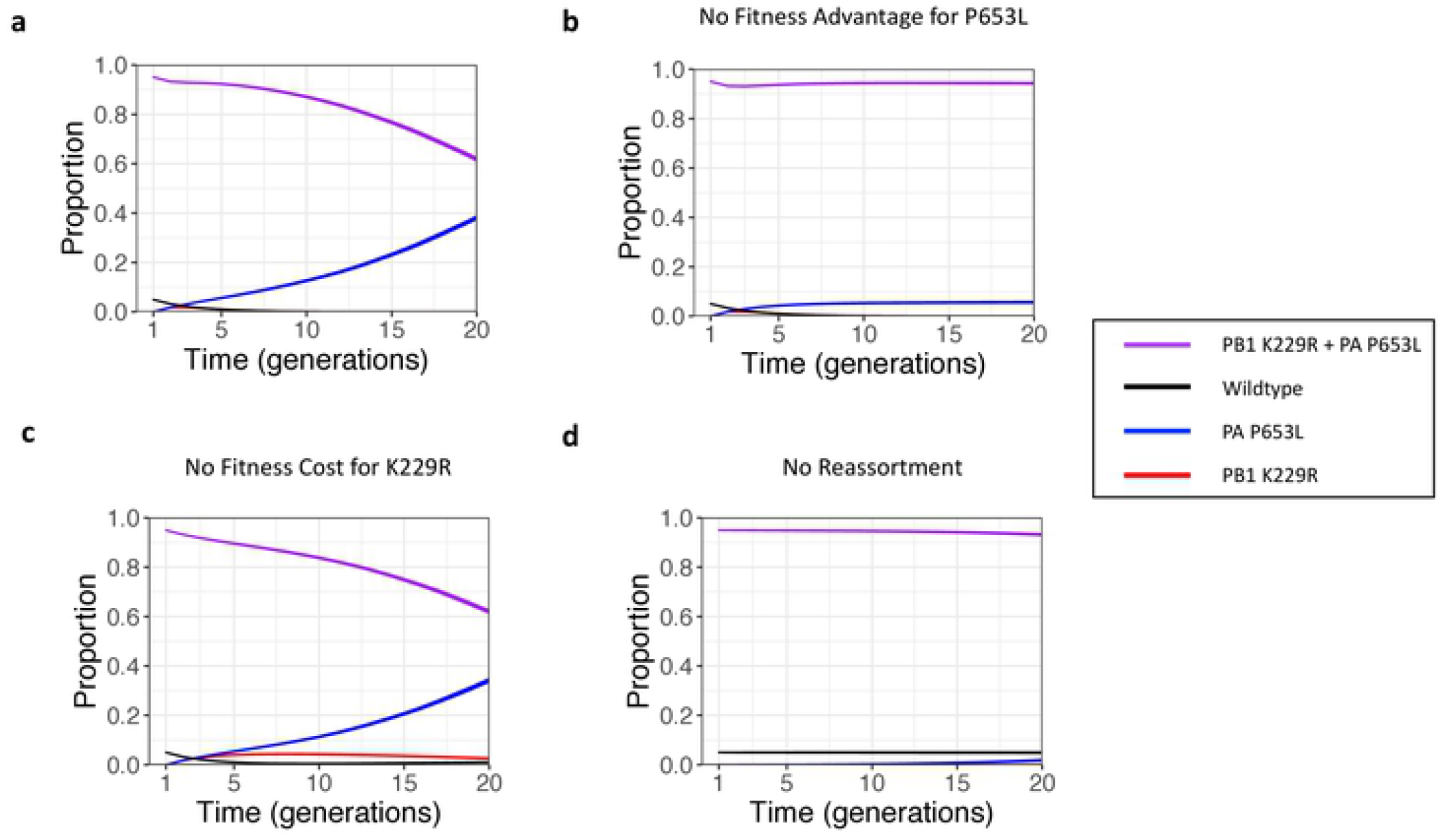
**a)** The proportion of each virus genotype are shown over 20 rounds of replication for a model with reassortment and mutation. The starting proportions are 5% Wild type and 95% K229R + P653L. Strain fitness for Wild type, K229R, P653L and K229R + P653L were set at 1, 0.01, 1.25 and 1 respectively. 10^6 viruses are modelled with 10^6 cells with a mutation rate, μ = 2 ×10^−4^. **b)** As **a** but the strain fitness for Wild type, K229R, P653L and K229R + P653L were set at 1, 0.01, 1 and 1 respectively. **c)** As **a** but the strain fitness for Wild type, K229R, P653L and K229R + P653L were set at 1, 1, 1.25 and 1 respectively. **d)** As **a** but there was no reassortment allowed during coinfection, only mutation. All graphs show results from 100 replicates (the line width is from the 2.5th to the 97.5^th^ percentile).

Next, we removed reassortment from the model allowing genotypes to change only through selection and mutation (Figure 3d). Without reassortment, there was very little change in genotype frequencies following 20 generations of the model implying that reassortment was necessary to accurately model the observed dynamics. This is because co-infection with wild type and K229R + P653L, and thus the opportunity for the P653L single mutant to be generated through reassortment, occurs much more frequently than de novo mutation for the MOI and mutation rates assumed. This model was robust to changes in values of the initial variables as shown in the sensitivity analysis, unless the mutation rate and/or the fitness of the PA P653L mutant were much larger (see Appendix).

### PA P653L does not show a fitness benefit in more recent viruses

Given the large fitness advantage of the P653L mutant virus in ferrets, it is surprising that the mutation has not been observed in pH1N1 isolates. We hypothesized that one reason the mutation might not be present is that more recent mutations in the pH1N1 virus polymerase also confer a fitness advantage, achieving the same increase in polymerase activity as P653L. Our previous work showed that the polymerase of Eng195 was less well adapted to human cells compared to later pandemic isolates and identified the N321K PA mutation as being a key mutation that led to improved polymerase activity in second and third wave pH1N1 virus isolates(29, 31). To test whether there was epistasis between N321K and P653L, we introduced the N321K mutation into Eng195 and tested polymerase using the minigenome activity (Figure 4a). Eng195 PA N321K had higher polymerase activity compared to P653L (1-way ANOVA, p<0.001). There was no difference in polymerase activity between N321K and N321K + P653L (1-way ANOVA, p=0.80). This implied that N321K provides a greater increase in polymerase activity than P653L and P653L provided no additional benefit to polymerase activity in the presence of N321K.

**Figure 4.**
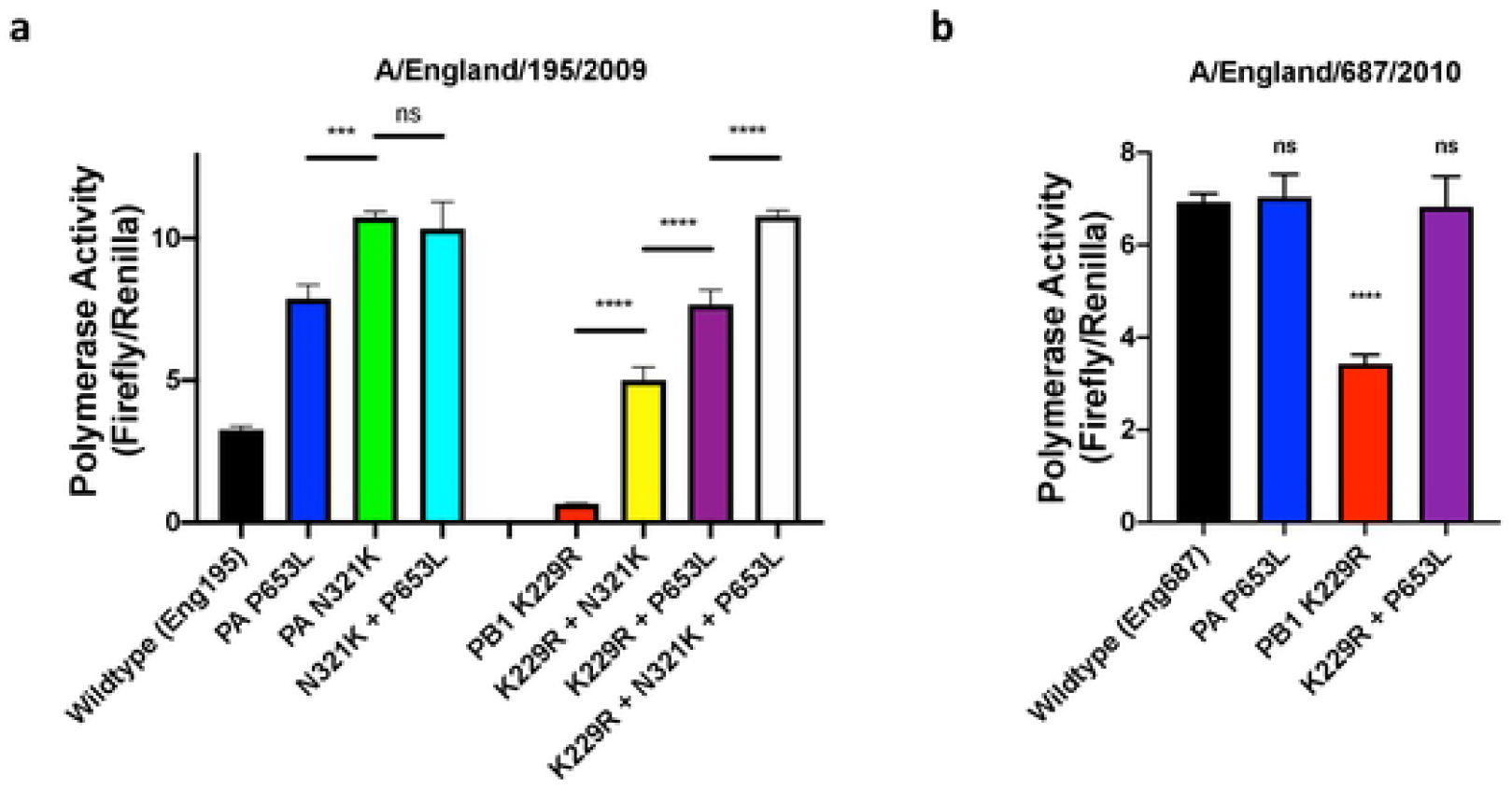
Minigenome assays were performed in 293T cells. Pol I –firefly luciferase minigenome reporter, at 0.08 μg and PCAGGS-Renilla, at 0.1 μg were transfected with PCAGGS plasmids coding for wildtype and mutated polymerase subunits (PB1, PB2 and PA) and NP at 0.08, 0.08, 0.04 and 0.12 μg respectively derived from **a)** Eng195 first wave and **b)** Eng687 third wave pH1N1 virus. Luciferase signal was read 24 hours post-transfection. Polymerase activity is given as a ratio Firefly to Renilla signals. One-way ANOVA with Dunnett’s multiple comparison test, *** p<0.001, **** p<0.0001, ns= not significant.

Next, we wanted to test whether N321K could affect the evolution of resistance of favipiravir by compensating for the defect in polymerase activity caused by PB1 K229R. Introducing both K229R and N321K into Eng195 showed that N321K partially compensated for the loss of polymerase activity conferred by K229R but did not reach the level of polymerase activity of P653L + K229R (Figure 4a). Adding the compensatory mutation, P653L to K229R + N321K showed full compensation to the level of N321K. Next we introduced the K229R and P653L mutations into the polymerase of a representative 3^rd^ wave pandemic H1N1 virus, Eng687, which already contained N321K. As had been observed in the background of Eng195, Eng687 K229R resulted in low but appreciable polymerase activity which was fully compensated by the presence of P653L (Figure 4b). However, cost to polymerase activity caused by K229R was noticeably less in Eng687 than in Eng195.

## Discussion

In this study, we showed that favipiravir-resistant influenza A virus could productively infect ferrets and transmit through contact transmission and via respiratory droplets. By infecting ferrets with a mix of wild-type and favipiravir resistant viruses, we sought to determine whether there was a fitness difference between the viruses. Although, the PB1 R229 and PA L653 mutations were initially present on two RNA segments within the same virus, their evolutionary trajectories were decoupled over time in the ferrets. In individual ferrets where there was a mix of K229 and R229, there was an increase in the wild-type PB1 K229 residue over time. Our modelling showed that this was not due to the fitness cost of the R229 mutation as that was adequately compensated by L653, but rather due to a fitness advantage for the P653L single mutant. This fitness advantage was implied in our previous work where the single PA mutation conferred higher polymerase activity in a minigenome assay as well as a slight growth advantage in the virus at 24 hours in cell culture(17). Despite the fitness advantage of the P653L single mutant, there was a slight increase in frequency of P653 within some ferrets. (19)In the donor ferrets, two ferrets showed an increase and two, a decrease in the proportion of P653. We suggest that this was likely due to stochastic changes in the proportion of wild-type viruses to mutant viruses within the donor ferrets. No ferret had a frequency of P653 higher than K229 confirming that the main driver of frequency changes was the fitness advantage of the P653L single mutant. P653 did not transmit to any of the sentinel ferrets almost certainly due to low frequency of P653 and the small bottleneck size of transmission. Whilst our simple model was capable of showing the rise in the P653L single mutant, it could not reproduce the stochastic changes in the frequency of P653. A spatially explicit model might have the additional complexity necessary to model these stochastic dynamics.

It is notable that in 4/5 ferrets where only R229 transmitted, K229 was not generated by reversion of the K229R mutation demonstrating that reassortment was necessary for the initial production of the K229 + L653 virus and that the R229 + L653 virus was fit and capable of a productive infection. The one exception was the group 3 ferret infected through indirect contact, which showed no K229 at the first sequencing time point but 38% K229 at the second sequencing time point. Whilst this could have been caused due to reversion to K from R229, it could have also been due to subsequent reinfection from the donor (animals were exposed to donors throughout the experiment) or by outgrowth of low levels of K229, which were initially present but were not detected by sequencing. Our modelling supports that the virus with single PA mutation L653 coupled with wild-type PB1 K229 was generated by reassortment in vivo before increasing in frequency due to positive selection (Figure 5). The high levels of coinfection and reassortment necessary for such a scenario have been previously seen in experimental infections of ferrets and other animals(32, 33).

**Figure 5.**
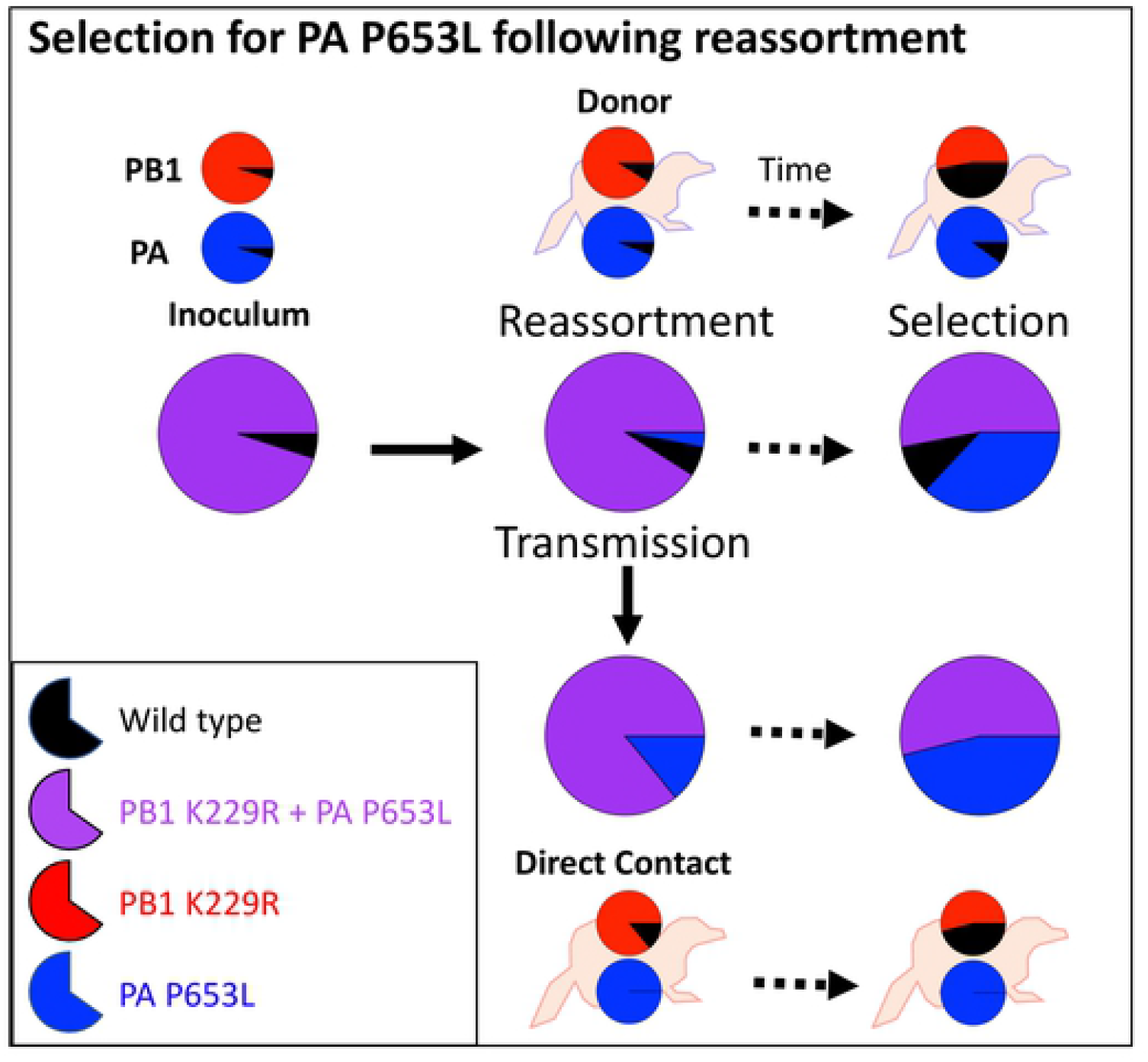
Schematic explaining how virus populations change for the donor and direct contact ferrets from Group 1. Large pie charts show the percentage of PB1 K229R + PA P653L mutant (purple) and wild-type viruses (black). Reassortment leads to the generation of the single mutant PA P653L (blue) in the donor which is transmitted to the direct contact. The proportion of PA P653L increases over time due to positive selection. Wildtype virus did not transmit and increases (or decreases) stochastically over time in donor ferrets. Smaller pie charts on each ferret show the sequencing results for PB1 and PA as in Figure 2.

Despite the large fitness advantage of L653, this mutation is not found in currently circulating pH1N1 viruses. We showed that one potential explanation for this observation is that other PA mutations have evolved that ameliorate the maladapted PA of the early pH1N1 isolates compared to more recent pH1N1 isolates. We have previously shown that later isolates from the third wave could outcompete first wave isolates in vitro due to the N321K mutation in PA(29). Here we showed that Eng195 polymerases reconstituted with PA harbouring N321K derive no additional fitness benefit from L653. This is the most likely explanation for why L653 has not been seen in sequenced isolates. Surprisingly, PA N321K could partially compensate for the low polymerase activity of PB1 K229R despite not being structurally close to the active site. This might imply that favipiravir-resistance could evolve more easily in more recent pH1N1 viruses as the essential resistance mutation in PB1 does not suffer such a dramatic fitness cost. However, in a polymerase constellation based on the third wave virus Eng687 harbouring K229R in PB1, the P653L mutation was still required to fully compensate and to attain comparative activity to the wild-type polymerase.

Our results have interesting implications for transmission dynamics of favipiravir resistant viruses in the absence of drug. We have demonstrated that the virus can transmit between ferrets that are untreated with drug, which means that a localised epidemic of drug resistant virus would be possible. However, we also found that the K229R mutation was lost over time within a single ferret which implies that if the virus can be outcompeted, resistance is less likely to spread in the absence of drug. The exact probability of resistance emerging and resistance then spreading will likely depend on the precise genetic background of the virus. For example, although the K229R resistance mutation was lost over time in our experiment based on a first wave pH1N1 2009 virus, K229R might not have been lost, at least by reassortment with wild-type virus, in more modern pH1N1 viruses where there is no benefit to polymerase activity of the compensatory mutation P653L in the absence of K229R. In the event of a new pandemic the specific fitness effects of both resistance mutations as well as of compensatory mutations will determine the likelihood of drug resistance emerging and spreading. Given that the compensatory mutation, P653L, has not been found in sequenced isolates, it suggests that robust resistance will always require multiple mutations. However, if resistance does arise, resistant viruses might continue to spread locally in the absence of drug pressure in a permissive genetic background.

## Materials and Methods

### Cells and Virus

Madin-Darby canine kidney (MDCK; ATCC) and HEK293T (293T) were grown in Dulbecco’s modified Eagle’s medium (DMEM; Invitrogen) supplemented with 10% fetal bovine serum (FBS; labtech.com), 1% penicillin-streptomycin (Invitrogen) and 1% non-essential amino acids (Gibco) at 37 °C and 5% CO2.

A/England/195/2009 (Eng195) is a first-wave isolate from the 2009 A(H1N1) pandemic grown from a reverse genetic virus(29). Favipiravir resistant Eng195 virus (K229R + P653L) containing a K229R mutation in PB1 and a P653L mutation in PA was constructed as described previously(17).

### P653L proportions in sequenced viruses

pH1N1 (2009) viruses with full length PA segments were downloaded from GISAID. They were aligned to an Eng195 reference and all mutants at location 653 were analysed in Geneious.

### Animal Studies

Female ferrets (20–24 weeks old) weighing 750–1000 g were acclimatized for 14 days before inoculation. Donor ferrets were lightly anaesthetized with ketamine (22 mg/kg) and xylazine (0.9 mg/kg) and then inoculated intranasally with virus diluted in phosphate buffered saline (PBS) (0.1 ml per nostril). The virus inoculum was ∼10,000 plaque forming units consisting of a mix of K229R + P653L virus and Eng195 in the ratio 95:5. Sentinel ferrets were introduced day 1 post infection and remained for the duration of the experiment. Ferret body weight was measured daily to check for significant weight loss due to sickness. Ferrets were nasal washed daily, while conscious, by instilling 2 ml PBS into the nostrils, and the expectorate was collected in 250 ml centrifuge tubes. Virus titre in the nasal wash expectorate was calculated by plaque assay. The nasal wash was stored with 4% Bovine Serum Albumin Factor V (Gibco) at −80 °C prior to RNA extraction. All sentinel ferrets were handled before donor ferrets to prevent accidental transmission of virus. All animal research described in this study was approved and carried out under a United Kingdom Home Office License, PPL 70/7501 in accordance with the approved guidelines.

### Sequencing

Samples were chosen at multiple time points for each ferret (Figure 1) and sequenced at Public Health England. Viral RNA was extracted using easyMAG (bioMérieux) and one step Reverse-Transcription-PCR was performed with Superscript III (Invitrogen), Platinum Taq HiFi Polymerase (Thermo Fisher) and influenza specific primers. To ensure coverage of PB1 and PA, gene specific primers were used to amplify PB1 and PA which were sequenced in parallel with the whole genome samples. Sequencing libraries were prepared using Nextera library preparation kit (Illumina) and sequenced on an Illumina MiSeq generating 150-bp paired end reads. Reads were mapped with BWA v0.7.5 and converted to BAM files using SAMTools (1.1.2). Variants were called using QuasiBAM, an in-house script at Public Health England. Raw sequences have been deposited at https://www.ebi.ac.uk/ena (project number PRJEB39934.)

### Modelling

Viral evolution was modelled over the time using an individual-based model. The model tracked the proportion of free virions of the wild-type virus Eng195, the resistant virus K229R + P653L and viruses containing a single mutation, K229R or P653L. The model was initialised with 10^6^ virions, a mix of K229R + P653L virus and Eng195 in the ratio 95:5, as in the inoculum, and assumes a well-mixed population with virions infecting cells randomly. Evolution of the virus was tracked over 20 generations of replication, with 10^6^ cells available for infection during each generation giving an average MOI of 1. Both the number of virions and the population of cells stayed constant between generations. We allowed for varying amounts of reassortment by adjusting the ratio of viruses to cells. During each replication cycle, each virion enters a random cell. The burst size from each cell was Poisson distributed, with mean proportional to the summed fitnesses of the virion(s) infecting that cell. In the baseline model, Eng195, K229R, P653L and K229R + P653L were assigned a relative fitness, with default values equal to 1, 0.01, 1.25 and 1 respectively. Within a cell, new segments were produced with a probability equal to the proportion of founding virions in that cell. These segments were randomly combined into virions, allowing reassortment, assuming each virus RNA segment in the cell had an equal chance, unrelated to genotype, of being incorporated into progeny virus. Newly produced virions mutated either PB1 or PA with probability μ = 2 ×10^−4 (1)^. Excess virions were discarded randomly to maintain the viral population size.

For the model with mutation only, each cell produced whole virions for each strain, proportional to the founding virions for each strain in that cell. The newly produced virions were mutated as per above.

Code to reproduce model results can be found at https://github.com/ada-w-yan/reassortment/.

### Minigenome Assay

To measure polymerase activity, pCAGGS plasmids containing genes encoding PB1, PB2, PA and NP from Eng195 or A/England/687/2010 (Eng687) were transfected into 293T cells using Lipofectamine 3000 (Thermo Fisher). Plasmids containing the K229R PB1, N321K PA and P653L PA mutations were constructed by site-directed mutagenesis. Plasmid quantities per well were PB1-0.08 µg, PB2-0.08 µg, PA-0.04 µg and NP-0.12 µg. In addition, a PolI-luc plasmid (0.08 µg), encoding a firefly luciferase minigenome reporter with influenza A segment 8 promoter sequences, was transfected with a pCAGGS-*Renilla* luciferase control (0.1 µg). After 21 hours, cells were lysed and luciferase activity measured using the Dual-Luciferase Reporter Assay kit (Promega). Polymerase activity was expressed as the ratio of Firefly:*Renilla*. Polymerase combinations were compared using 1-way ANOVA with p-values adjusted using Dunnett’s Multiple Comparison test.

## Figure Legends

**S1-Appendix**. This appendix details the sensitivity analysis for the modelling described in Figure 3.

